# Dynamic Monomer-Dimer Transition in Ligand-induced Apelin Receptor Activation

**DOI:** 10.1101/2025.02.19.639188

**Authors:** Su-Yu Ji, Wei-Wei Wang, Yixin Yang, Ping Xu, Jiangrong Zhang, Xinyue Zhao, Kun Xi, Shao-Kun Zang, Dan-Dan Shen, Chunyou Mao, Qingya Shen, Yan Zhang

**Author notes:** These authors contributed equally: Su-Yu Ji, Wei-Wei Wang, Yixin Yang.

## Abstract

G-protein-coupled receptors (GPCRs) are significant signal transducers that exist as monomers and in multiple oligomeric forms. However, molecular mechanism driving their dynamic interconversion to regulate intricate signaling in class A GPCRs remains elusive, compounding our understanding of their related pathophysiological functions. Here, we present a set of 12 assemblies of the apelin receptor (APLNR), including dimeric apo state, agonistic small molecule- or nanobody-bound state of monomeric and dimeric APLNR with and without G-proteins, providing a detailed dynamic view of the monomer-dimer transition. High-resolution cryo-EM structures reveal that different ligands induce varying degrees of pre-dissociation of dimers in the absence of G-protein, with G-protein coupling facilitating the transition from dimeric to monomeric receptor. Functional studies further highlight the critical role of cholesterol clusters in stabilizing the APLNR dimers. These insights enhance our understanding of the dynamic regulation of class A GPCRs across different aggregated forms and advance the rational drug design strategies aimed at selectively modulating of APLNR signaling.

## Introduction

GPCRs serve as central hubs in cellular signal transduction, ensuring the complexity and precision of downstream signaling through their structural versatility, characterized by diverse aggregated forms and dynamic conformational changes^1-4^. While Class C GPCRs are obligate dimers for their function, Class A GPCRs exist in a dynamic equilibrium between monomeric and oligomeric forms^5-10^, exhibiting distinct agonist affinity, efficacy, as well as the signaling profiles and pharmacological functions^11-14^. However, it remains unclear whether the various aggregated forms mediate signaling transduction in an interconvertible manner or independently (Fig. 1a), compounding our understanding of complex mechanism of downstream signaling. Further studies on the molecular basis and functional correlations of various aggregated forms and their interconversion in class A GPCRs are essential for achieving precise modulation of downstream signaling and understanding their pathophysiologic roles.

**Fig. 1.**
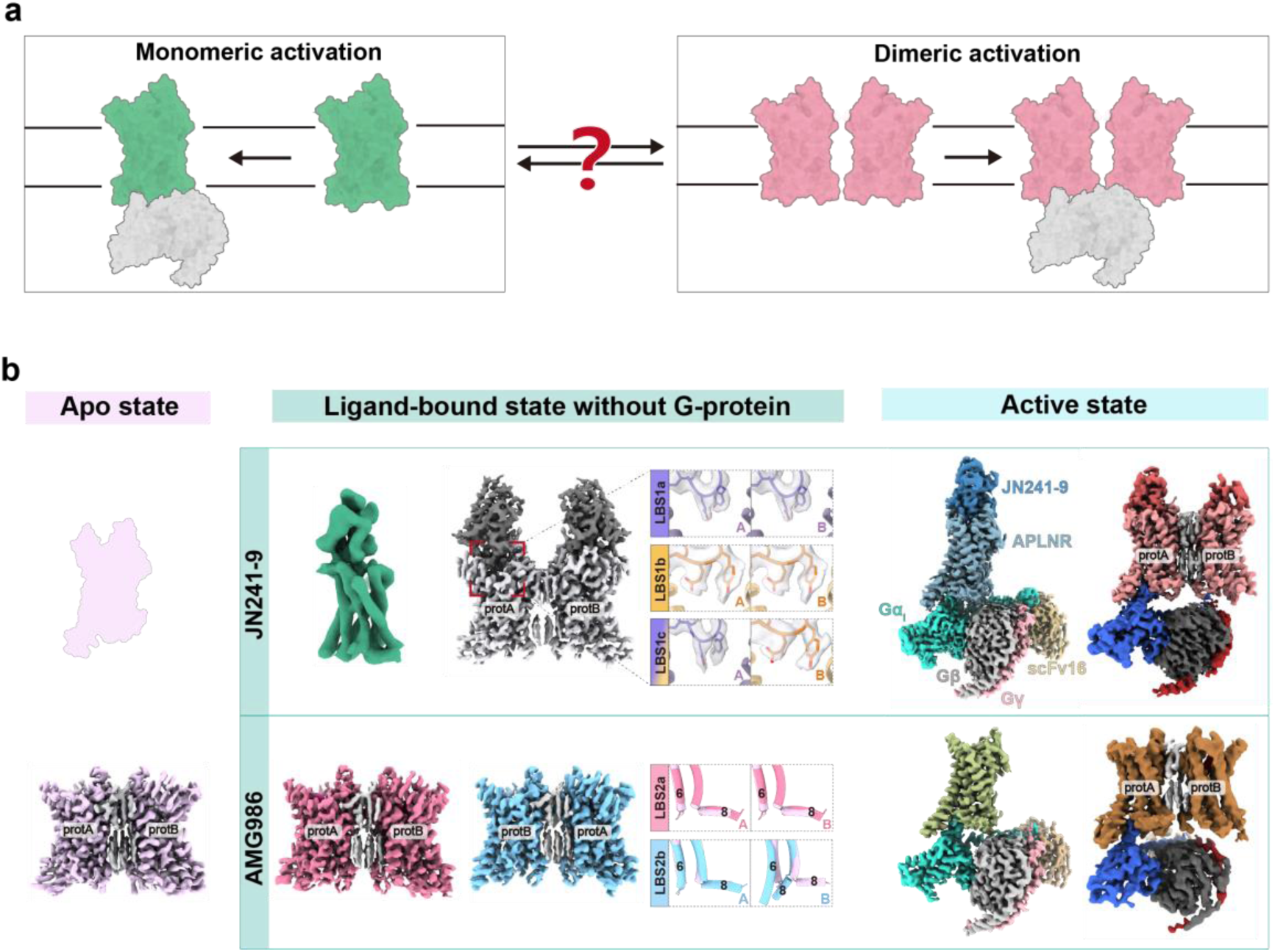
Cryo-EM structures of APLNR monomers and dimers bound to different ligands during the activation process. **a,** Schematic representation of whether the monomeric and dimeric activation processes of class A GPCR are independent or interconvertible. **b,** Cryo-EM density map of APLNR across various conformational states. The plum planar cartoon illustration represents the apo-state monomer, which exists but is challenging to capture; Apo-state dimer: plum; JN241-9–monomer: sea green; JN241-9-bound APLNR dimer without G-protein contains three conformations that differ only in the ligand, in which the receptor adopts the same inactive conformation, represented by gray. Three distinct conformations of JN241-9 include symmetric LBS1a: slate blue, LBS1b: orange, and asymmetric LBS1c: a mixture of slate blue and orange; AMG986-bound state of APLNR dimers: LBS2a is shown in pale violet red and LBS2b is shown in sky-blue; For the JN241-9–monomer–G_i_ complex: JN241-9: steel blue, APLNR: light slate gray, Gαi: turquoise, Gβ:silver, Gγ: light pink, scFv16: tan; For the JN241-9–dimer–G_i_ complex: JN241-9: indian red, APLNR: light indian red, Gαi: royal blue, Gβ: dim gray, Gγ: brown, scFv16: tan; For the AMG986–monomer–G_i_ complex: AMG986: dark olive green, APLNR: light olive green, Gαi: turquoise, Gβ: silver, Gγ: light pink, scFv16: tan; For the AMG986–dimer–G_i_ complex: AMG986: maroon, APLNR: deep yellow, Gαi: royal blue, Gβ: dim gray, Gγ: brown, scFv16: tan; Cholesterol molecules at the dimeric interface: light gray. The two protomers are designated as protA and protB, respectively.

APLNR stands out as an ideal model system for investigating the dynamic transition between monomeric and dimeric forms of class A GPCR, supported by single-molecule imaging that confirms dimeric formation, and it is the first Class A GPCR whose stable dimeric structures have been captured by cryo-EM^8,15,16^. However, these structures are inadequate for fully capturing the dynamics of the dimer-monomer transition, given their limited resolution and insufficient determination of multiple conformations. Moreover, the relationship between different aggregated forms and pathophysiologic functions has not been elucidated yet. APLNR is regard as a promising target due to its critical role in maintaining cardiovascular health and preventing muscle weakness^17-22^. Nonetheless, apelin-induced β-arrestin-dependent signaling contributes to detrimental cardiac hypertrophy, driving interests in the development of G-protein-biased APLNR agonists as potential heart failure therapies^23-26^. It is suggested that APLNR dimers exhibit distinct pathophysiological functions compared to monomers, indicating differences in the selective signaling pathways associated with various aggregated forms^27-30^. Therefore, it is crucial to elucidate how diverse ligands recognition induces different aggregated forms, ultimately impacting signaling and therapeutic effects. AMG986, also named as BGE-105 and Azelaprag, is the most potent small molecule agonist in activating both G-protein and β-arrestin signaling of APLNR, advancing rapidly in clinical trials for anti-obesity treatment^31-33^. However, a Phase 2 clinical trial evaluating AMG986 was recently discontinued owing to potential adverse effects, highlighting the necessity to explore its molecular mechanism in stimulating various apelin receptor forms. In parallel, JN241-9 stands as the pioneering single-domain antibody agonist targeting class A GPCR, derived from JN241. The greater size of nanobody agonist contributes to the alignment of particles in cryo-EM data processing, overcoming micelle interference, thereby facilitating the capture of monomeric APLNR in the absence of G-protein. We have selected these two agonists to conduct a comprehensive study on the activation mechanisms and functions of dimers and monomers.

In this study, we resolved 12 cryo-EM density maps of wild type full-length APLNR, encompassing monomeric and dimeric forms of the apo state, agonists-bound state in the presence or absence of G_i_-proteins, 9 of which achieved high resolution between 2.4 and 3.0 Å and enabled the accurate structure modelling. We therefore directly observed the stepwise transition and conformational changes of APLNR dimers towards monomers from the apo to the active state. Furthermore, combining with functional experiments, we elucidated the pivotal role of cholesterol clusters in stabilizing APLNR dimers, as well as how different ligands modulate the dimer-to-monomer transition, providing insights into drug development targeting different aggregated forms of class A GPCRs.

## Results

### Signaling and structural determination of APLNR dimers and monomers during activation process

Previous structural studies of the APLNR have indicated that the π-π interactions network formed by F101^3.24^ plays a central role in stabilizing the dimeric interface^15^. To investigate the differences in downstream signaling mediated by distinct aggregated forms, we first assessed the signaling profile of wild-type APLNR and the dimer-reduced form (F101^3.24^A) using AMG986 and JN241-9, by conducting a NanoBiT assay and a GloSensor cAMP assay to detect G-protein activity and the bioluminescence resonance energy transfer (BRET) assay to detect β-arrestin recruitment. AMG986 showed roughly 10-fold greater potency in activating various G-proteins, including G_i_, G_q_ and G_13_, as well as β-arrestin, compared to JN241-9 (Supplementary Figs. 1a-f). Additionally, AMG986 induced a more rapid activation rate than JN241-9 (Supplementary Fig. 1d). Though negligible differences in EC50 or Emax from G-protein signaling, the dimer-reduced form (F101^3.24^A) showed a higher Emax than the dimeric form in the β-arrestin signaling (Supplementary Fig. 1f), consistent with previous results^15^, indicating that the dimer exhibits G-protein bias compared to the monomer. Moreover, cAMP accumulation results showed that the dimer-reduced form exhibits higher constitutive activity than that of dimeric form (Supplementary Figs. 1d-e), prompting us to investigate the regulatory mechanism of dimerization induced by different potent ligands.

Determination of cryo-EM structures for class A GPCRs without G-protein is impeded by their small dimensions and inherent instability. Moreover, the crystallization process is rendered challenging when the structures exhibit flexibility, particularly when intermediate-state GPCRs manifest a diversity of conformations. Typically, receptors are often fused with proteins like BRIL to increase stability, which may not accurately represent their native conditions^34,35^. Here, we successfully achieved high-resolution cryo-EM structures of agonists-bound class A GPCR in various conformations without G-protein, delineating the dynamic process of activation. Specifically, dimeric receptors with increased size contributes significantly to achieve high-resolution structures in the absence of G-protein (Fig. 1b and Supplementary Figs. 1g-j). Furthermore, to overcome the difficulty of obtaining monomeric GPCR structures hindered by micelle interference, we selected JN241-9, a single-domain antibody, as an agonist suitable for structural alignment. Fortunately, we identified an obvious JN241-9–APLNR monomer species during 2D and 3D classification, albeit at relatively low resolution. Data processing statistics revealed a 5:1 ratio of dimer to monomer, with three distinct conformations of the JN241-9–dimer identified, differing only in the ligand-binding pose (termed by ligand-bound states LBS1a, LBS1b and LBS1c, respectively) (Fig. 1b and Supplementary Fig. 2a). Additionally, the two AMG986-bound states of APLNR dimer without G-protein are distinguished by unique conformational changes of TM6 and helix 8 (H8) in one protomer (termed by LBS2a and LBS2b, respectively) (Supplementary Fig. 2b). These results indicate that the conformations are in a ligand-dependent manner. Furthermore, we determined the structures of the APLNR–G_i1_ complexes with the two agonists. Data processing statistics clearly demonstrate that monomeric complexes greatly exceed dimeric complexes, though dimeric formation is preferred in the absence of G-protein (Supplementary Figs. 2 and 3), implying that most dimers have dissociated during activation. Notably, the two protomers are designated as protA and protB, respectively (Fig. 1b). Data processing revealed that AMG986-bound dimeric APLNR–G_i1_ complexes adopt two conformations in protB (termed by ACT2a and ACT2b, respectively), aligning with those structures without G-protein (Supplementary Fig. 3b).

### Ligands recognition and conformational comparison in different states

To elucidate the structural transition of APLNR from its apo to active state, we first compared two AMG986-bound states of APLNR dimer without G-protein (LBS2a and LBS2b) against the apo state to identify structural differences (Fig. 2a). In both apo and AMG986–dimer^LBS2a^ structures, the receptor adopts an inactive conformation, characterized by TM6 remaining in the closed state. In contrast to LBS2a, protB in LBS2b adopts an active-like conformation, with TM6 undergoing an outward movement and H8 bending inward to engage the intracellular pocket, in line with the previously reported fully active conformation (Fig. 2a)^15^. Our observations further reveal that protB progressively moves away from protA across the transition states from apo to LBS2b, represented by the movement of TM3 and TM4 in the intracellular half of protB when aligned with protA (Fig. 2b). While hydrophobic interactions involving the “FGTFF” motif (APLNR residues 97-101) contribute to dimeric stabilization in the extracellular half (Supplementary Fig. 4a), the intracellular half of the dimer interface is primarily stabilized by more extensive interactions mediated by cholesterol clusters, particularly involving TM1, TM2, and TM4 (Supplementary Fig. 4b). Cα distance measurements between TM1–TM4 residues of protB and nearby protA residues at the intracellular interface revealed a gradual increase from the apo state to LBS2b, with the most notable change at F78^2.53^ (Fig. 2c and Supplementary Figs. 4c-f). These results suggest that in the absence of G-protein, APLNR dimers bound to ligands such as AMG986 exhibit a tendency to dissociate into monomers.

**Fig. 2.**
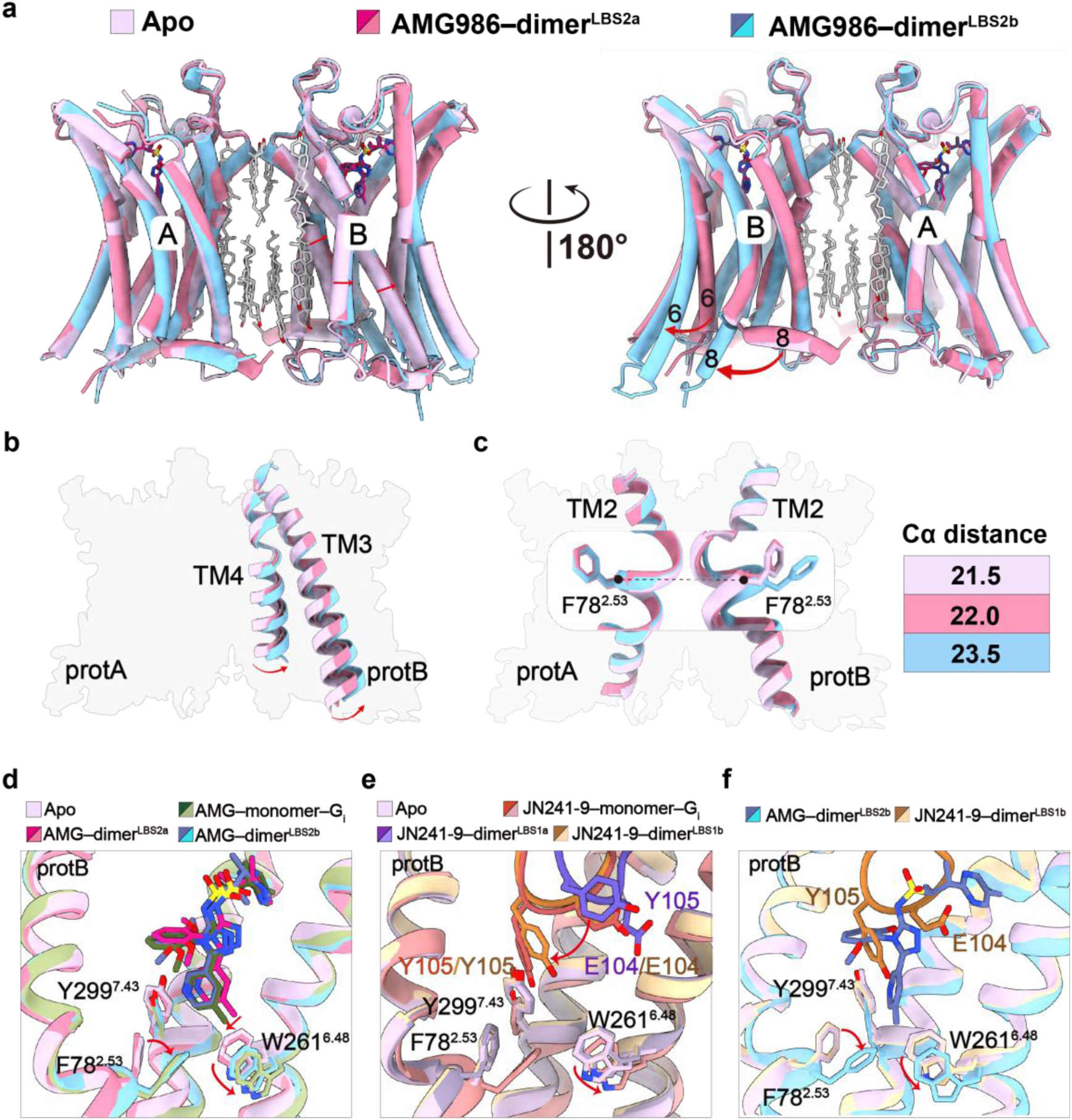
Conformational changes upon agonists binding to APLNR. **a,** Structural alignment of apo-state APLNR–dimer, AMG986-bound state of APLNR dimer without G-protein LBS2a and LBS2b. In symmetric LBS2a, protB adopts an inactive conformation, whereas in asymmetric LBS2b, protB adopts an active conformation with the outward displacement of TM6, and the helix 8 bends inward to occupy the intracellular pocket. **b,** ProtB is observed to gradually move away from protA when aligning with protA from the apo state to LBS2a and then to LBS2b, represented by TM3 and TM4. **c,** Utilizing Cα of residue F78^2.53^ as a reference point to quantify the distances between ProtA and ProtB across the apo state, AMG986-bound state LBS2a and LBS2b. **d,** Conformation changes of the binding pocket of apo state and AMG986-bound APLNR–dimer^LBS2a^ and dimer^LBS2b^, as well as AMG986-bound APLNR monomer–G_i_ complex. **e,** Conformation changes of the binding pocket of apo state and JN241-9-bound APLNR–dimer^LBS1a^ and dimer^LBS1b^, as well as JN241-9-bound APLNR monomer–G_i_ complex. **f,** Comparison of the binding pocket of AMG986-bound APLNR–dimer^LBS2b^ and JN241-9-bound APLNR–dimer^LBS1b^.

Upon comparing the ligand-binding pockets of the AMG986–dimer (LBS2a and LBS2b) and the AMG986–monomer–G_i_ complex, we noted that AMG986 in the protB of LBS2b adopts a similar binding pose to that in the AMG986–monomer–G_i_ complex (Fig. 2d). AMG986 in protB of LBS2b displays a lateral shift toward Y299^7.43^ relative to its position in LBS2a, thereby triggering a downward shift of key residues associated with activation, including F78^2.53^, W261^6.48^ and Y299^7.43^, ultimately resulting the outward movement of TM6 in protB of LBS2b (Fig. 2d). Surprisingly, we observed three distinct conformations of JN241-9 as aforementioned, while the receptor remained unchanged in its inactive conformation. The pivotal residue Y105, located in the CDR3 region of nanobody JN241-9 and inserting into the very bottom of the ligand pocket, undergoes a rotation from pointing towards the crevice between TM4/5 and ECL2 of APLNR in the LBS1a conformation to facing the intracellular direction in the LBS1b conformation (Figs. 1b and 2e). When comparing the ligand-binding pockets of JN241-9–dimer with and without G-protein, we noted that in LBS1b, the Y105 of JN241-9 is positioned closer to Y299^7.43^, resembling the fully active conformation. However, no alterations in key motifs or TM6 conformational changes were observed in LBS1b (Fig. 2e). In the fully active-state structure of JN241-9, Y105 exhibits a deeper insertion toward Y299^7.43^, facilitating the shift of F78^2.53^, W261^6.48^ and Y299^7.43^ into their activation conformations. Furthermore, comparison of the ligand-binding pockets in protB showed that AMG986 penetrates more deeply than JN241-9, thereby driving the allosteric transition of F78^2.53^, W261^6.48^ and Y299^7.43^, which could explain its capacity to induce the opening of TM6 (Fig. 2f). These findings indicate that AMG986 can independently induce activation of a single protomer within the APLNR dimer, in contrast to JN241-9, which necessitates G-protein engagement for receptor activation. We also observed that the ligand engagement induced conformational changes of key motifs in receptors across the entire activation process, including the rearrangement of the “Na^+^ pocket”, “PVF motif”, “NPxxY motif”, “DRY motif”, which align with the known mechanisms of GPCR activation (Supplementary Fig. 5).

### G-protein coupled activation of APLNR facilitates the dimer-to-monomer transition

According to statistics and analysis of cryo-EM data processing, while two agonists-bound APLNR in the absence of G-protein exhibits more dimers than monomers, the proportion of monomer–G-protein complexes in the fully active state far exceeds dimer–G-protein complexes, indicating that most of dimers dissociate into monomers when introducing G-protein (Fig. 3a, Supplementary Figs. 2 and 3). The protB in JN241-9–dimer–G_i_ complex adopts an inactive-state configuration and H8 in protB is disordered. By comparing the contact areas between the receptor and the G-protein in monomeric complex and dimeric complex, we found that the buried surface area of monomer (1213 Å^2^) is significantly greater than that of dimer (917.7 Å^2^), suggesting that G-protein binding to the monomer is more stable (Fig. 3b). In addition, we compared JN241-9- and AMG986-bound dimer^LBSb^ to their monomeric complex aligning with protA, and observed that the binding of heterotrimeric G-protein would generate a clash with protB in the dimer^LBSb^, therefore prompting the dissociation of protB (Fig. 3c).

**Fig. 3.**
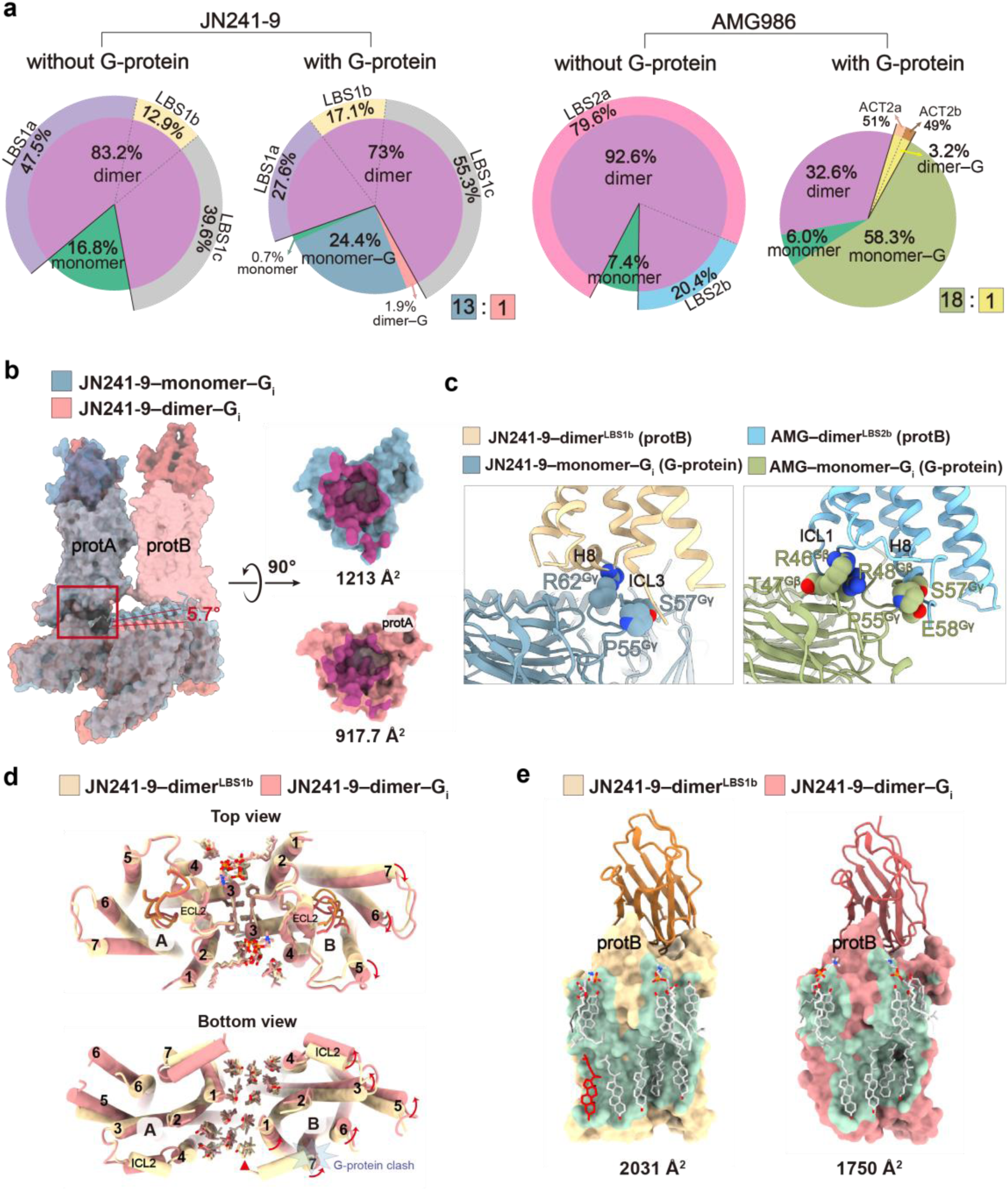
The binding of G-protein triggers the transition of APLNR from dimers to monomeric complexes. **a,** Cryo-EM data processing statistics and analysis of different conformations APLNR structures bound to JN241-9 and AMG986 in the presence or absence of G-protein, shown by pie charts. **b,** Comparison of the surface area on the receptor interacting with G-protein between JN241-9-bound monomeric and dimeric complexes. **c,** Details of the clash generated between the G-protein of the monomeric complex and the protB of the G_i_-free APLNR dimer bound to JN241-9 and AMG986. **d,** Comparison of the JN241-9–dimer–G_i_ complex and JN241-9–dimer^LBS1b^ aligning with protA from the top view and the bottom view. **e,** Surface representation of the interface between the cholesterol molecules and protB, with the buried surface area of 2031 Å^2^ in JN241-9–dimer^LBS1b^ and 1750 Å^2^ in JN241-9–dimer–G_i_ complex, respectively.

Meanwhile, there are still a small fraction of cryo-EM particles corresponding to dimer–G-protein complexes, accounting for 1/14 of monomeric complexes activated by JN241-9 and 1/19 of monomeric complexes activated by AMG986 (Fig. 3a). The high-resolution structure of the JN241-9–dimer–G_i_ complex shows a 5.7-degree downward rotation of G-protein measured by αN helix to allow for the stable binding of protB. Notably, the position of the αN helix in the JN241-9–dimer–G_i_ complex closely resembles that observed in the apelin–dimer–G_i_ complex (Fig. 3b and Supplementary Fig. 7e). Additionally, cholesterol clusters at the dimerization interface were observed to elastically connect the two protomers like an adhesive sponge (Fig. 3d). Compared to the JN241-9-dimer^LBS1b^, the JN241-9–dimer–G_i_ complex exhibits a counterclockwise rotation of protB based on TM4 from the bottom view, which moves away from protA (Fig. 3d). The cholesterol density near TM1 is disorder in dimeric complex, resulting in a decreased protB-cholesterol interacting area of 1750 Å^2^ in the dimeric complex compared to that of 2031 Å^2^ in the JN241-9–dimer^LBS1b^, which suggest that protB tends to dissociate from protA when bound to G-protein (Fig. 3e). Furthermore, the kinetic GloSensor cAMP assay revealed that signaling response associated with monomeric activation (F101A) was greater than that of the dimeric form (WT) during the initial 15 minutes. However, both responses converged thereafter, further suggesting that most dimers transition into monomers upon receptor activation.

Careful cryo-EM data processing of the JN241-9-bound APLNR–G_i1_ complex identifies a class of dimers that were not bound to G-protein, from which we further uncover three distinct fractions of particles, whose conformations are consistent with the G_i_-free specimens. Intriguingly, the particle number ratios of these conformations change from 47.5% (LBS1a), 12.9% (LBS1b) and 39.6% (LBS1c) to 27.6%, 17.1% and 55.3%, respectively. The increasement of the proportions of LBS1b and LBS1c states further indicates that these two conformations are closer to the active state conformation in comparison with LBS1a state (Fig. 3a, Supplementary Figs. 2a and 3a).

### The extent of dimer dissociation upon APLNR activation is ligand-dependent

Different from JN241-9, the sole action of AMG986 without G-protein is capable of achieving the partial activation of one protomer in the APLNR dimer, thus prompting a tendency for the protomers to dissociate from each other (Fig. 2 and Supplementary Fig. 4). Inspiringly, the structures of AMG986–dimer–G_i_ complexes adopt either the active or the inactive conformations in protB, aligning with those dimeric structures in the absence of G-protein (Supplementary Fig. 6a). These findings suggest that ligand-specific modulation of dimeric conformations may contributes to varying degrees of dimer dissociation upon activation.

Comparison of the AMG986–dimer^LBS2b^ with the monomeric complex reveal that the binding of G-protein would generate a more severe clash between Gβγ in the monomeric complex and H8 as well as ICL1 of protB in the AMG986–dimer^LBS2b^, compared to the JN241-9-bound APLNR structures (Fig. 3c). This implies that the AMG986-bound dimer has a higher propensity to dissociate during activation than the JN241-9-bound dimer. Additionally, comparing the two agonists-bound dimeric complexes highlights a counterclockwise rotation of about 27° in the entire G_i_ heterotrimer observed in the AMG986-bound structure (Supplementary Fig. 6b).

To date, multiple agonist-bound APLNR dimer structures in the presence or absence of G_i_-protein have been resolved. Accordingly, we compared the APLNR structures resolved in this study with previously reported structures. Pairwise alignment of the entire dimeric assemblies revealed relatively small Cα root mean square deviation (RMSD) values (<1.0 Å) among G-protein-free dimers (Supplementary Fig. 7a). In contrast, Cα RMSDs between structures with and without G-protein coupling exhibited larger deviations (1.0 to 2.5 Å), largely attributable to conformational shifts in TM6 and H8 (Supplementary Fig. 7a). Moreover, Cα RMSDs among G-protein-coupling dimers also showed relatively larger deviations (1.0 to 2.1 Å), suggesting that different ligands induce distinct dimer conformations in the active state (Supplementary Fig. 7a). To assess the extent of dimer dissociation, we compared the Cα RMSDs of protB aligned on protA (Supplementary Fig. 7b). Comparison between G-protein-bound and G-protein-free dimers revealed a remarkably wide range of Cα RMSDs (2.3 to 6.3 Å), particularly in comparisons involving the ELA–G_i_ complex (PDB: 7W0N), further indicating different ligands induce varying degrees of dimer dissociation during activation (Supplementary Figs. 7b-d).

### Cholesterol plays an important role in stabilizing APLNR dimerization

The fluidity of the phospholipid bilayer permits membrane proteins dispersed therein to achieve sufficient lateral mobility, meanwhile, cholesterol promotes membrane fluidity and stiffness by interweaving into the hydrophobic gaps between the phospholipid acyl chains^36,37^. Cholesterol is typically enriched in lipid rafts and plays an important role in stabilizing the specific conformations of membrane proteins, ensuring their proper functions. Previous studies on cholesterol–GPCR interactions have focused on class A and class B GPCR–G-protein complex^38-40^, as well as class C GPCR dimers^41^, with limited research on class A GPCR dimers owing to the lack of related structures. Our density maps show that at the dimeric interface of APLNR, two parallel rows of cholesterol molecules are located at the intracellular half to stabilize dimeric formation, while fewer cholesterols are present on the extracellular half, attributed to the stability of hydrophobic interactions (Figs. 4a-b). Remarkably, well-defined densities with the shape of phospholipid in the extracellular half of the dimeric interface were observed in all dimeric structures (Supplementary Fig. 8a). In addition, we also observed phospholipids in the extracellular half between TM5 and TM6 in the APLNR dimeric structure bound by JN241-9 (Supplementary Fig. 8a). These suggest that the phospholipids in the dimeric structure also play a role in stabilizing dimerization.

**Fig. 4.**
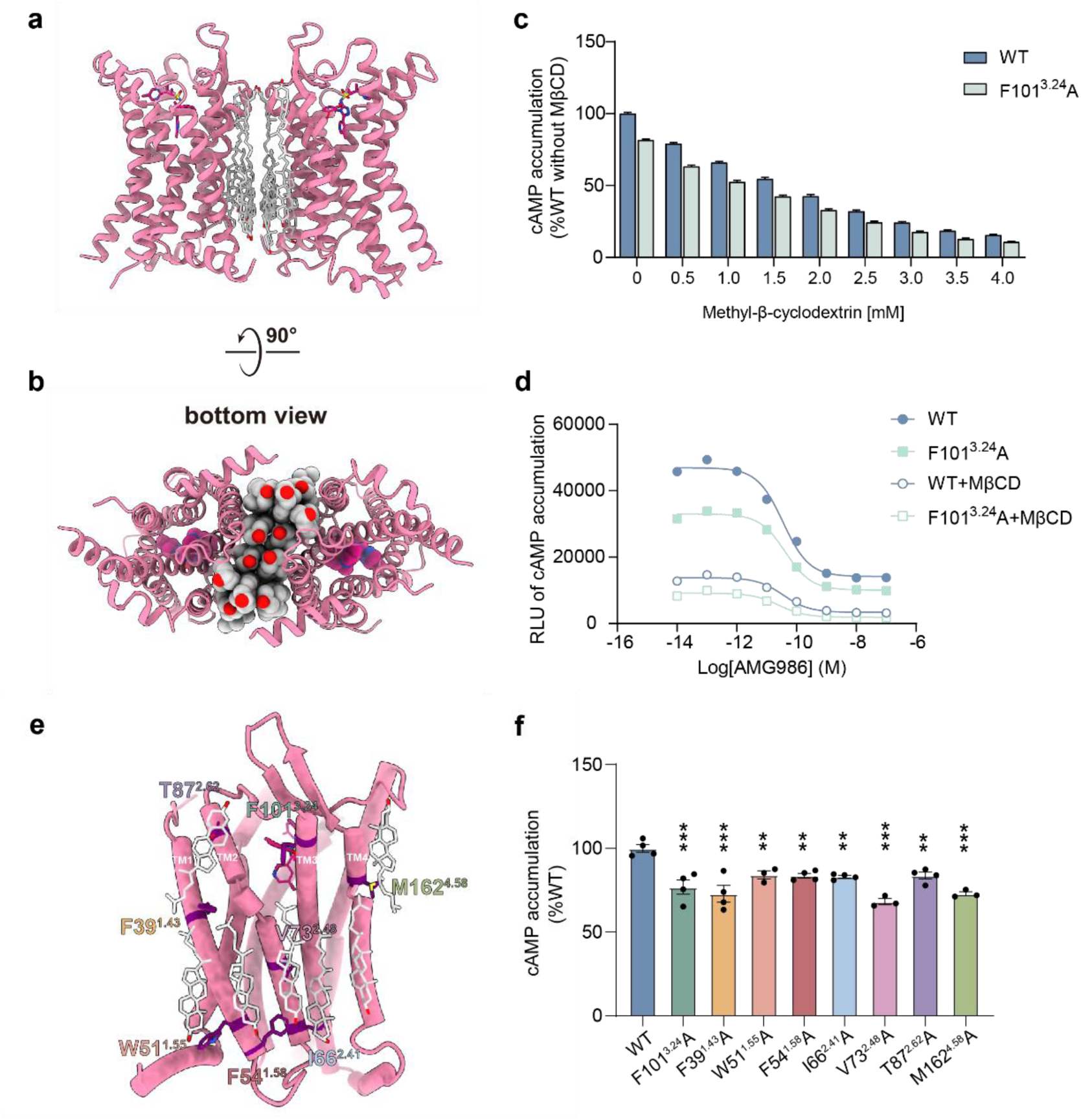
Cholesterol clusters in dimeric interface play a significant role in stabilizing dimeric formation. **a, b,** Side view (**a**) and bottom view (**b**) of the APLNR dimeric interface. Cholesterol clusters located at the dimeric interface are depicted in light gray. **c,** Histogram depicting the effect of cholesterols removal by Methyl-β-cyclodextrin at various concentration on the basal activity of cAMP accumulation in WT and F101^3.24^A mutant, as measured by GloSensor cAMP assay in HEK293T cells. **d,** Dose-response curves for AMG986-induced cAMP accumulation in WT and F101^3.24^A, with or without treatment with 3 mM cyclodextrins, as measured by the GloSensor cAMP assay in HEK293T cells. **e,** Display of amino acid residues in dimeric interface interacting with cholesterols. **f,** Measurement of basal activity in WT and single-point mutants of residues interacting with cholesterols using the GloSensor cAMP assay in HEK293T cells.

To investigate the stabilizing effect of cholesterol on dimerization, we detected cAMP accumulation after treating with Methyl-β-cyclodextrin (MβCD) at various concentrations to deplete cholesterols from the cell membrane. We found that depletion of cholesterol led to a decrease in the value of cAMP accumulation without affecting the expression of receptors (Fig. 4c and Supplementary Fig. 8b). This decrease reflects an enhancement in basal activity, indicating that the cholesterol depletion promotes the dissociation of dimers into monomers. In addition, F101^3.24^A exhibits higher basal activity than WT at every concentration of the MβCD, indicating that the hydrophobic interaction formed by F101^3.24^ and cholesterols between the dimer play a role in stabilizing the dimer together (Fig. 4c). The cAMP accumulation curve measured at 3mM MβCD further suggests that MβCD addition did not influence EC50 and Emax (Fig. 4d). Furthermore, we mutated the key residues that interact with cholesterol in the dimerization interface to small residues Ala or Gly, and found that the basal activity increased, further illustrating the importance of cholesterol in stabilizing receptor dimerization (Figs. 4e-f and Supplementary Fig. 8c).

## Discussion

The various aggregated forms of class A GPCRs trigger complex signaling networks, yet the mechanisms of interconversion between these forms remains unclear, hindering our comprehension of their related pathophysiological roles^4,11-14^. In this study, we have resolved 12 monomeric and dimeric structures of APLNR across diverse states. These results elucidate that various ligands induce varying degrees of pre-dissociation of dimers without G-protein, a process further enhanced by G-protein coupling, which promotes the transition from dimers to monomers. Combining with functional assays, we further revealed the crucial role of cholesterol clusters in stabilizing the dimeric form of APLNR.

Previous studies indicate that the ratio of dimers and oligomers increases with receptor density on the cellular membrane^8,9^, particularly within lipid rafts. Additionally, certain pathological conditions are often accompanied by the overexpression of receptors and the formation of oligomers^30,42-45^. Comprehensive cryo-EM analysis of purified APLNR receptors revealed that APLNR predominantly exists as dimers in the absence of G-protein coupling. However, the binding of G-protein induces a clash with protomer B, facilitating its dissociation to form a more stable monomer–G-protein complex. Meanwhile, a minor population of dimer–G-protein complex was also captured in the fully active state, probably due to the concomitant rotation of the protB and G-protein, allowing their mutual accommodation (Fig. 5). Furthermore, our study suggests that different ligands induce diverse conformations during APLNR activation (Supplementary Fig. 6a). In the C2-symmetrical inactive LBS1a-1c and LBS2a complexes, G-protein binding to either protA or protB yields identical conformations (Scenario Ⅰ). For AMG986-bound LBS2b, G-protein binding to either protA or protB must overcome certain energy barriers. Binding to protB induces the assembly of the ACT2a complex (Scenario Ⅱ). In this instance, the α5 helix of Gα replaces the H8, driving protB from a partially active to a fully active conformation, while protA remains in its inactive state. On the other hand, the binding of G-protein to protA triggers the formation of the ACT2b complex (Scenario Ⅲ), wherein protA undergoes a classical activation mechanism characterized by an outward movement of TM6. Meanwhile, protB remains in a partially active state, with H8 bending inward to stabilize the open conformation of TM6, thereby preventing its transition to the fully active state (Scenario Ⅲ).

**Fig. 5.**
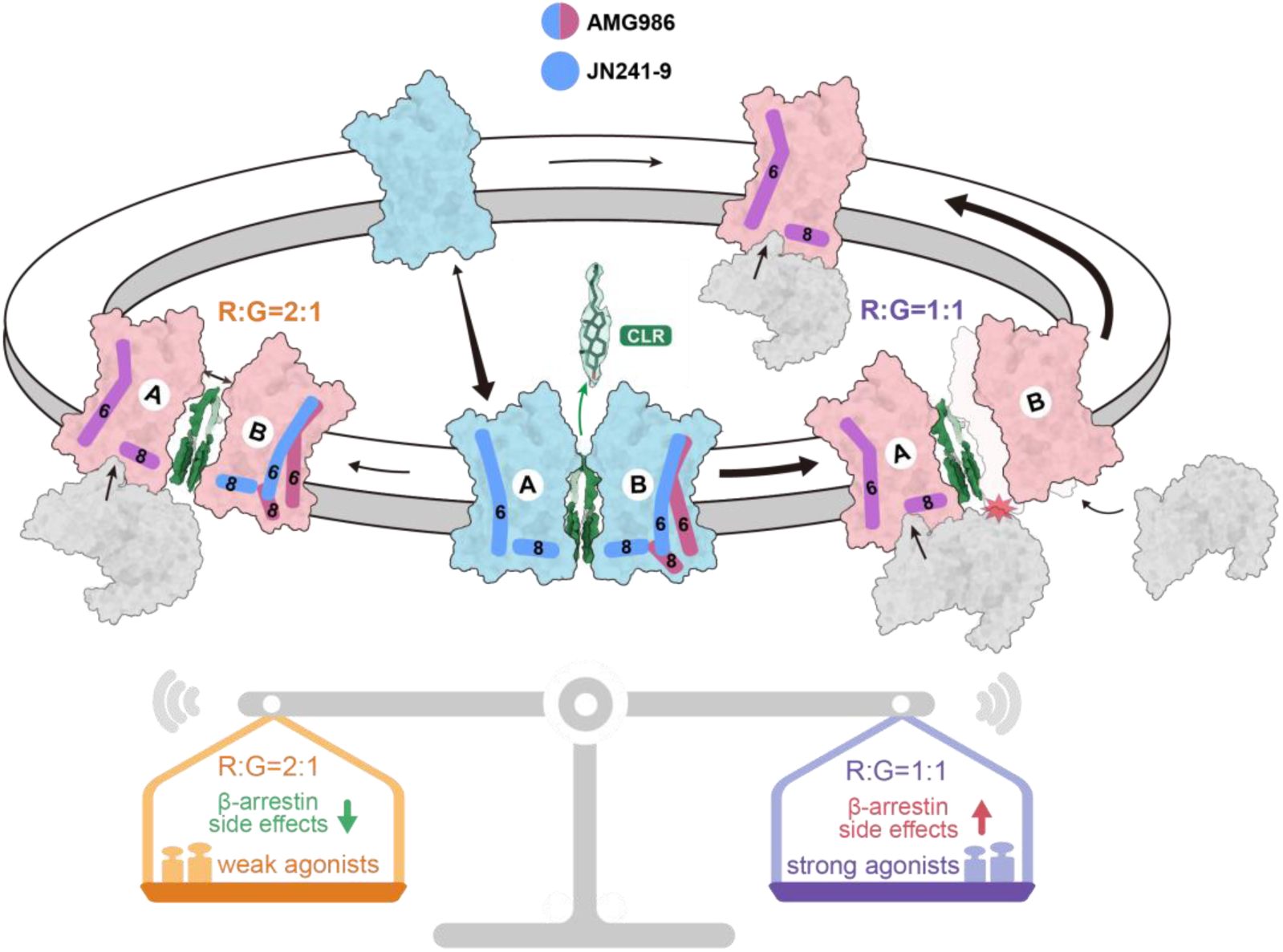
Schematic representation of the dynamic transitions between APLNR monomers and dimers during activation process. The schematic diagram illustrates that in the absence of G-protein coupling, monomers and dimers can interconvert with each other with dimers accounting for the majority. Both the monomer and dimer can transition to a monomer–G-protein complex upon agonists activating, and minority of dimers can also form a dimer–G-protein complex. The balance schematic diagram represents the downstream signaling bias of monomeric complex (R:G=1:1) and dimeric complex (R:G=2:1) induced by different ligands. CLR: cholesterols; R: receptor; G: G-protein; The black arrows indicate the interconversion between different conformations, with the thickness of the arrows representing the proportion of each conformation; The red star represents the clash between G-protein and protB.

Notably, although JN241-9 is insufficient to activate APLNR in the absence of G-protein, the transition of the ligand to the active conformation can still be captured (Fig. 2d), laying the foundation for the full activation of APLNR. Various G_i_-free conformations, induced by ligands with varying potency and different insertion depths into the binding pockets, lead to a distinct degree of pre-dissociation of protB (Fig. 2b and Supplementary Fig. 4), indicating that the dissociation from dimers to monomers during activation may be initiated and determined in the ligand-bound state of APLNR without G-protein coupling.

Cryo-EM data revealed a lower ratio of dimeric to monomeric complexes bound to AMG986 compared to JN241-9, suggesting that AMG986 more significantly promotes the formation of monomeric complexes than JN241-9. Notably, the monomeric APLNR exhibits greater β-arrestin activity than the dimeric form, while ligands-induced G-protein activity remains the same (Supplementary Fig. 1f), representing the increased potential of the receptor monomer to induce side effects of myocardial hypertrophy. These findings indicate that ligands characterized by higher potency and more profound insertion into the binding pocket tend to favor the formation of monomeric complexes, thereby enhancing β-arrestin signaling and consequently elevating the potential for adverse effects (Fig. 5). Inspiringly, solely prioritizing high potency and high affinity of ligands may not be optimal, as increased potency can also be accompanied by increased toxicity. A comprehensive consideration of various factors, including signaling differences arising from various aggregated forms, is essential for rational drug design. We also speculate that, though the eliminated β-arrestin activity of truly G-protein biased agonists WN353 and WN561 is primarily attributed to the altered interactions with “twin hotspots”, their reduced capacity to induce dimeric dissociation might also contribute to the weakening of β-arrestin signaling^23^.

Cholesterol is a critical component of cellular membranes, especially being abundant in lipid rafts where GPCRs are enriched and involved in the regulation of various biological processes^46,47^. Compared with phospholipids, cholesterols enhance the stiffness and order of the cell membrane surface, while also regulating membrane fluidity. It is considered indispensable for the structural stability and signaling regulation of various membrane proteins including GPCRs, and its position cannot be substituted by phospholipids^36,39,41,48-50^. In this study, we observed that cholesterol clusters were well arranged at the dimerization interface of APLNR, with the cholesterols-protomer contact area measuring 2031 Å^2^ in JN241-9–dimer^LBS1b^ (Fig. 3e), which is 14 times greater than the hydrophobic interaction interface (140 Å^2^). Additionally, in conjunction with functional studies, we provide a thorough understanding of how cholesterols modulate signaling pathways. These findings underscore the significance of considering the function and localization of lipids, such as cholesterols, in addition to protein-protein interactions, within the field of protein design, when predicting the intrinsic membrane protein interactome.

In summary, we have mapped out a detailed schema of the dynamic regulation between APLNR monomers and dimers using nine high-resolution cryo-EM structures, illustrating the complexity, diversity and tendency of dimeric activation modulated by ligands and cholesterols. Our work not only sheds light on the dynamic regulation of class A GPCRs in different aggregated forms, but also provides a structural basis for the precise design of drugs targeting diverse aggregated forms of APLNR aiming to reduce the on-target side effects.

## Acknowledgements

We thank the Cryo-Electron Microscopy Center of Liangzhu laboratory for help with cryo-EM data collection of apo, AMG986-, JN241-9-bound APLNR and AMG986-, JN241-9-bound APLNR–G_i1_–scFv16 complexes. This project was supported by the “Pioneer” and “Leading Goose” R&D Program of Zhejiang (2024C03147 to Y.Z.); National Natural Science Foundation of China grant (32430051, 92353303, 32141004, 81922071 to Y.Z., 32330049 to Q.S., and 32400575 to W.W.); the Key R&D Projects of Zhejiang Province (2021C03039 to Y.Z.); The STI2030-Major Projects (2022ZD0205400 to Q.S.), the China Postdoctoral Science Foundation (2024T170783 to W.W.); Postdoctoral Fellowship Program of CPSF (GZC20232326 to W.W.); Zhejiang Provincial Postdoctoral Research Project (ZJ2024043 to W.W.); Hubei Province Key Laboratory of Ischemic Cardiovascular Disease Open Fund Project (SZ202405 to W.W.); Y.Z. is also supported by the Fundamental Research Funds for the Central Universities and Peak Discipline Cultivation Program of Zhejiang University School of Basic Medical Sciences.

## Author contributions

Y.Z. initiated the study, conceived and supervised the whole project; Y.Z., Q.S., S.-Y.J., W.-W.W., and Y.Y. participated in the data analysis and interpretation. S.-Y.J., W.-W.W. designed the constructs of APLNR and expressed the proteins; S.-Y.J., W.-W.W. and P.X. purified these protein complexes; Y.Y. generated APLNR mutants for the cell-based G-protein activity assays and β-arrestin recruitment assays. Y.Y. generated APLNR mutants for the cell-based cAMP accumulation assays. X.Z. and D.-D.S. evaluated the samples by negative-stain EM; S.-K.Z. collected the cryo-EM data and S.-Y.J., performed cryo-EM data processing. S.-Y.J., Y.Y. performed model building; Y.Y. and J.Z. performed the cellular functional assays. S.-Y.J. and W.-W.W. performed structural analysis supervised by Y.Z.; W.-W.W., S.-Y.J. prepared the figures; S.-Y.J., W.- W.W., Q.S., and Y.Z. wrote the manuscript; Y.Z., Q.S., K.X. and C.M. provided important discussions and essential revisions. S.-Y.J., W.-W.W. and Y.Y. provided Figs. 1-5, Supplementary Figs. 1-8 and Tables 1-2.

## Notes

### Competing Interest Statement

The authors have declared no competing interest.

### Summary of Updates

Author affiliations updated. Supplemental files updated. We have revised the terminology used for the Gi-free dimer structures, replacing "INT" with "LBS" (Ligand-Bound State) to avoid any misinterpretation of these structures as intermediates. We also have replaced the term "intermediate-state dimer structures" with "dimers without G-protein" to enhance clarity. Additionally, we have added detailed structural comparisons between the previously reported APLNR structures and those resolved in this study. We have also revised the discussion section to more accurately describe the conformational changes of APLNR dimers upon G-protein binding.

