## Supplemental Figures 1-8 and Supplemental Tables 1-2 for "Dynamic Monomer-Dimer Transition in Ligand-induced Apelin Receptor Activation"

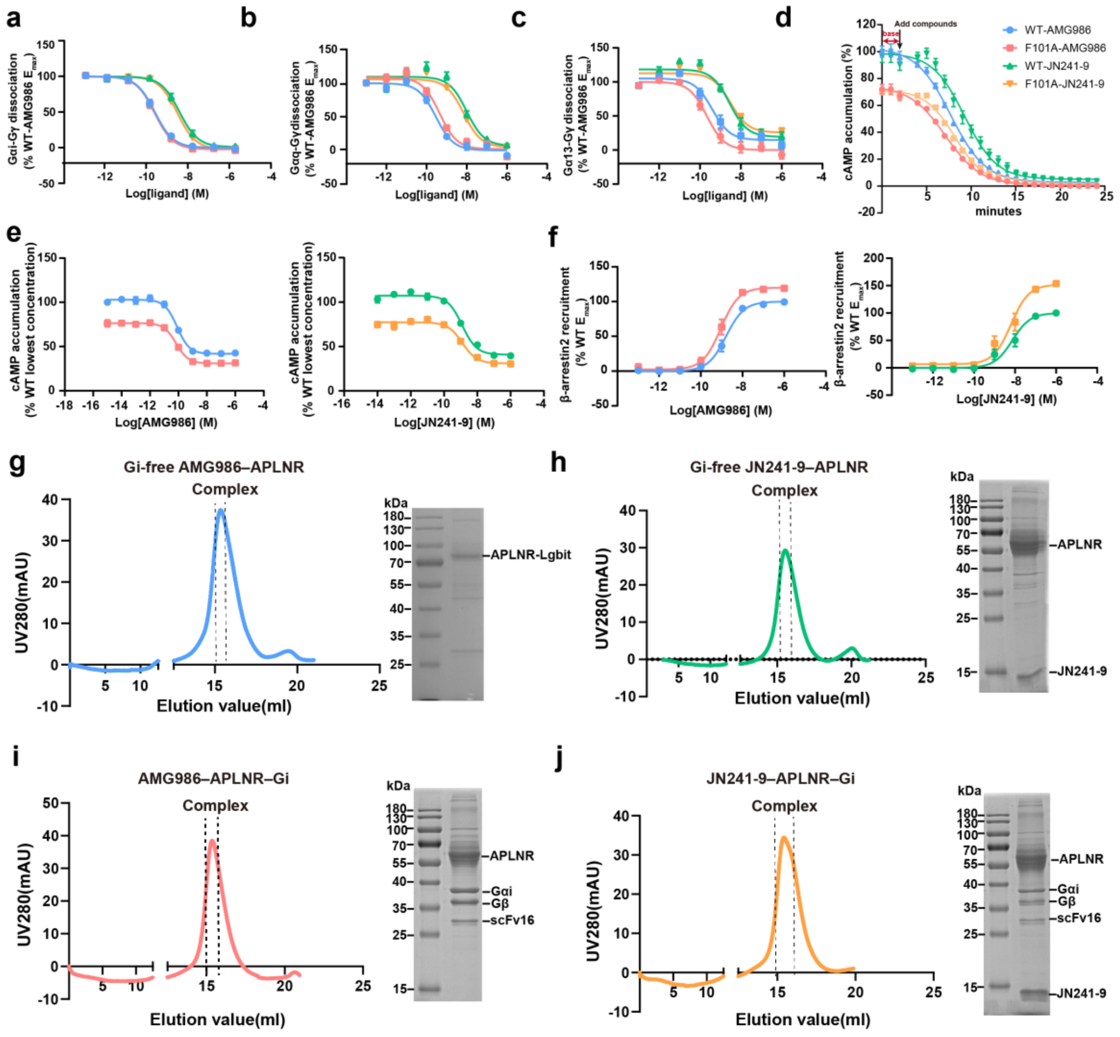

**Supplementary Figure 1. Analysis of ligands-triggered signaling pathways and purification of APLNR complexes in different states.**

**a-c,** The effects of two agonists on WT and partially dimer-disrupted mutant F101A. Dose-response curves show AMG986- and JN241-9-induced $G_i$ (**a**), $G_q$ (**b**) and $G_{13}$ (**c**) signaling were measured by NanoBiT assay in HEK293T cells.

**d,** Dose-response curves show the duration of cAMP signaling response induced by 10 μM of each ligand.

**e,** Dose-response curves for AMG986- and JN241-9-induced cAMP accumulation measured by GloSensor cAMP assay in HEK293T cells.

**f,** β-arrestin2 recruitment were measured by BRET assay.

**g-j,** Size-exclusion chromatography profile and SDS-PAGE gel of the purified $G_i$-free AMG986–APLNR complex (**g**), $G_i$-free JN241-9–APLNR complex (**h**), AMG986–APLNR–$G_i$ complex (**i**) and JN241-9–APLNR–$G_i$ complex (**j**).

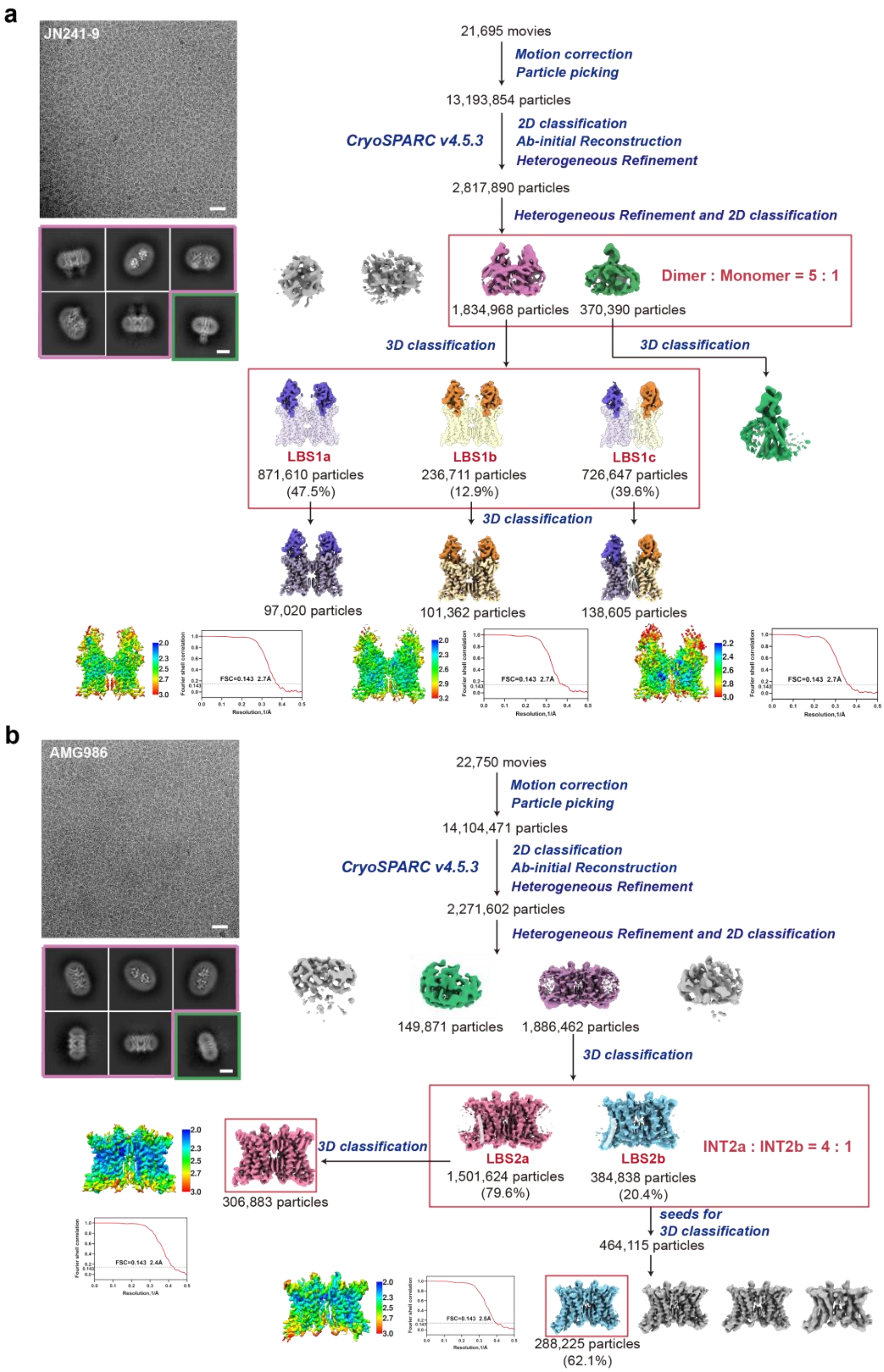


**Supplementary Figure 2. Cryo-EM data collection and processing for agonists-bound APLNR without G-protein coupling.**

**a, b,** Representative Cryo-EM micrographs (scale bar: 30 nm), representative reference-free 2D class averages (scale bar: 5 nm), flow chart of cryo-EM data processing, and 3D density map colored according to local resolution (Å) of the JN241-9- (**a**) and AMG986- (**b**) bound states of APLNR without G-protein coupling.

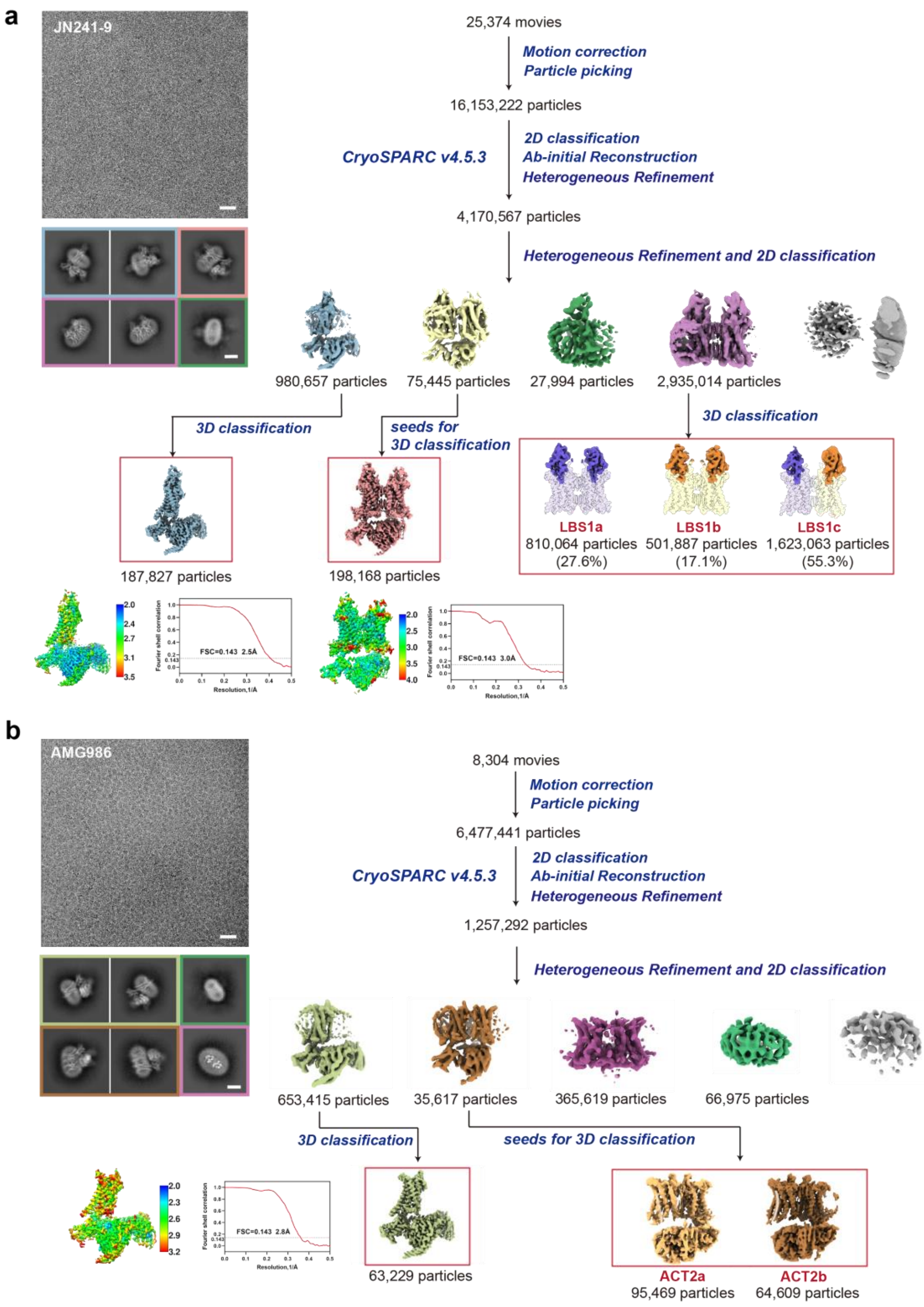


**Supplementary Figure 3. Cryo-EM data collection and processing for agonists-bound APLNR–$G_i$ complex.**

**a,b,** Representative Cryo-EM micrographs (scale bar: 30 nm), representative reference-free 2D class averages (scale bar: 5 nm), flow chart of cryo-EM data processing, and 3D density map colored according to local resolution (Å) of the JN241-9- (**a**) and AMG986- (**b**) bound APLNR–$G_i$ complexes.

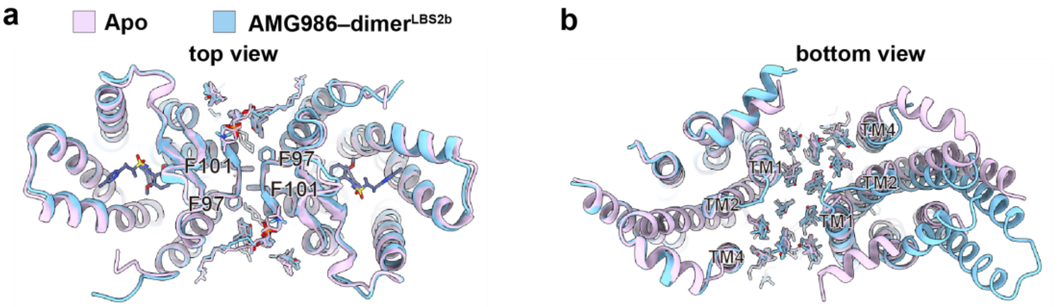


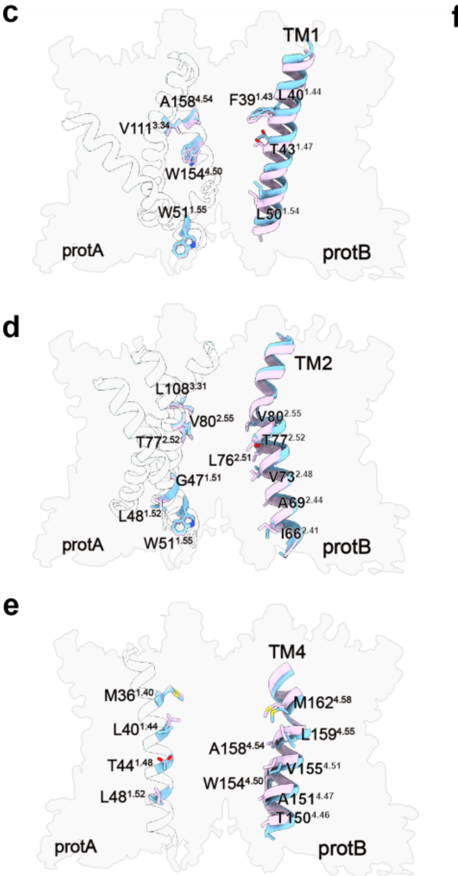


**f** Apo | AMG986–dimer$^{LBS2b}$

| Residue pairs | Cα distance (Å) Apo | Cα distance (Å) AMG986–dimer$^{LBS2b}$ |
|---|---|---|
| F39$^{1.43}$ — V111$^{3.34}$ | 23.9 | 24.9 |
| L40$^{1.44}$ — A158$^{4.54}$ | 20.1 | 21.3 |
| T43$^{1.47}$ — W154$^{4.50}$ | 20.6 | 21.4 |
| L50$^{1.54}$ — W51$^{1.55}$ | 16.0 | 17.1 |
| I66$^{2.41}$ — W51$^{1.55}$ | 18.6 | 19.1 |
| A69$^{2.44}$ — L48$^{1.52}$ | 20.7 | 21.4 |
| V73$^{2.48}$ — G47$^{1.51}$ | 17.0 | 18.0 |
| L76$^{2.51}$ — T77$^{2.52}$ | 17.1 | 18.5 |
| T77$^{2.52}$ — V80$^{2.55}$ | 14.8 | 16.4 |
| V80$^{2.55}$ — L108$^{3.31}$ | 17.1 | 18.4 |
| V111$^{3.34}$ — L76$^{2.51}$ | 23.7 | 24.1 |
| T150$^{4.46}$ — L48$^{1.52}$ | 21.4 | 22.4 |
| A151$^{4.47}$ — T44$^{1.48}$ | 19.0 | 20.0 |
| W154$^{4.50}$— T44$^{1.48}$ | 19.9 | 20.8 |
| V155$^{4.51}$ — T44$^{1.48}$ | 19.8 | 20.6 |
| A158$^{4.54}$ — L40$^{1.44}$ | 20.7 | 21.7 |
| L159$^{4.55}$ — L40$^{1.44}$ | 21.9 | 23 |
| M162$^{4.58}$— M36$^{1.40}$ | 22.6 | 23.4 |

**Supplementary Figure 4. Structural analysis of the dimeric interface and Cα distance measurement of residues within TM1-TM4 between protA and protB.**

**a-b**, Top view (**a**) and bottom view (**b**) of the APLNR dimeric interface in the apo state and AMG986-bound LBS2b.

**c-e**, ProtB moves away from protA in the AMG986-bound LBS2b relative to the apo state, represented by residues in TM1 (**c**), TM2 (**d**) and TM4 (**e**).

**f**, Cα distance measurement of residues within TM1-TM4 in different states. Each residue pair is presented with the first residue corresponding to protB and the second to protA, indicating their respective positions across the dimeric interface.

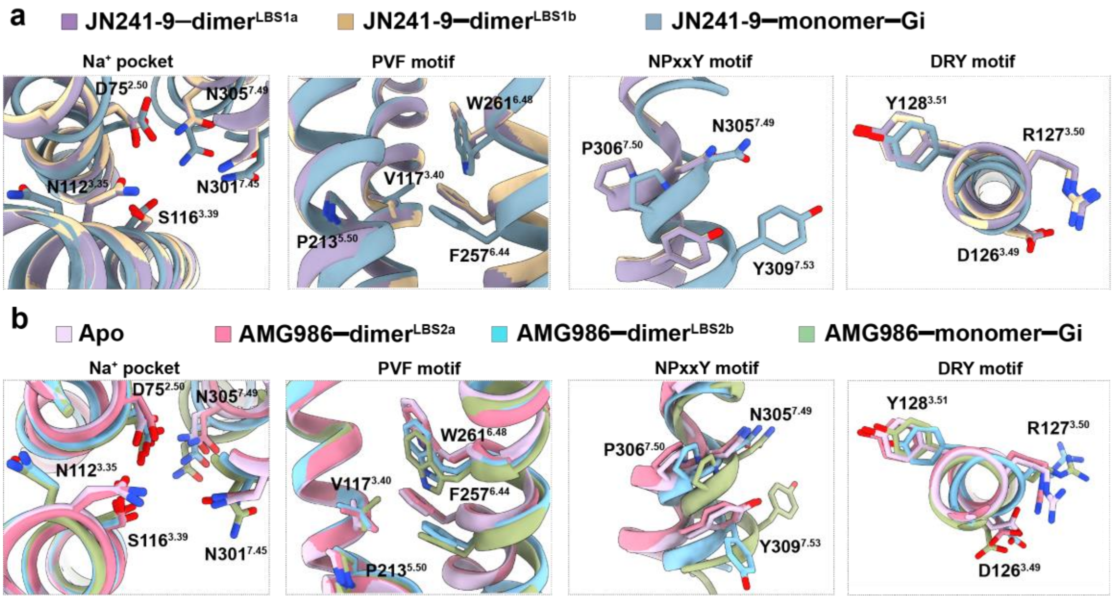


**Supplementary Figure 5. Activation mechanism of APLNR from the apo state to the active state.**

**a, b,** Conformational differences of important residues and motifs in APLNR activation induced by JN241-9 (**a**) and AMG986 (**b**), including “$Na^+$ pocket”, ‘‘PVF motif’’, NPxxY motif and “DRY motif”.

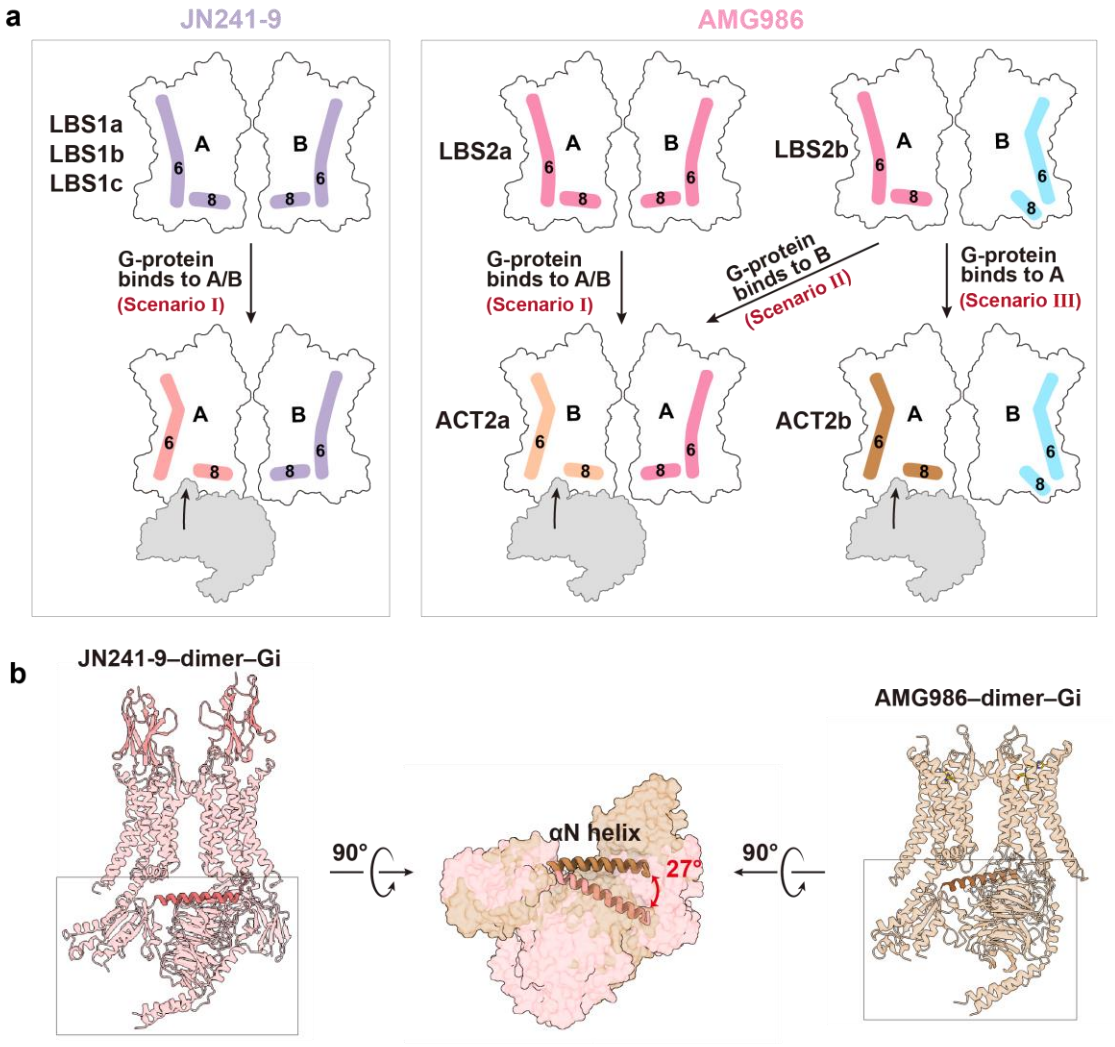


**Supplementary Figure 6. Conformational changes of APLNR dimerization are modulated by different agonists.**

**a,** Cartoon representations illustrate that G-protein binding to APLNR dimers with distinct conformations results in different dimer–G-protein complex conformations.

**b,** Comparison of the G-protein conformational changes represented by αN helix between the JN241-9- versus AMG986-bound APLNR dimer–$G_i$ complex.

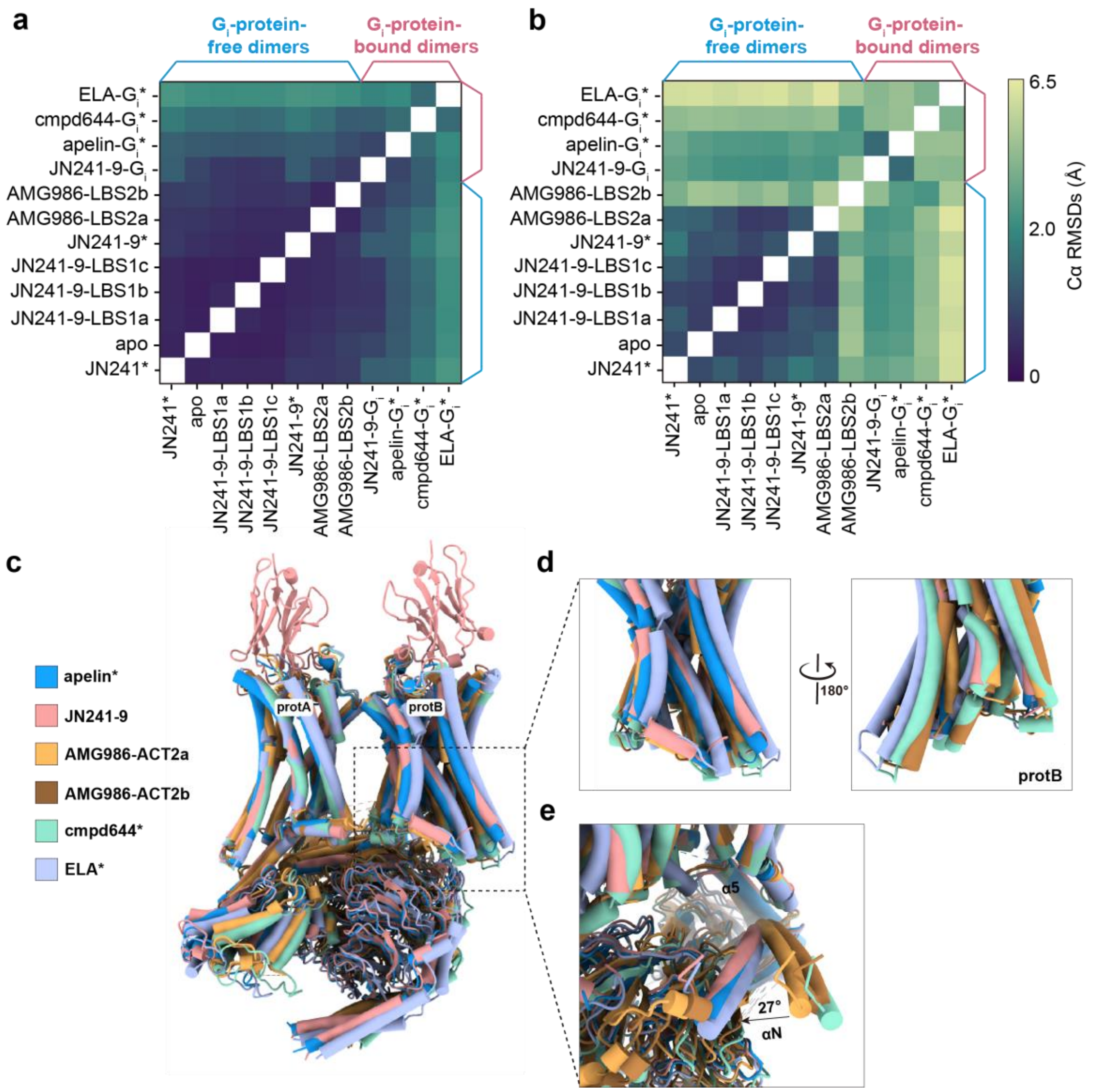


**Supplementary Figure 7. Structural comparison of previously reported and current dimeric APLNR structures in the presence or absence of G-protein.**

**a,** Heat map graphs display Cα RMSD values generated from pairwise alignments across the full APLNR dimers, assessing overall conformational differences.

**b,** Pairwise alignments were performed using protA of the APLNR dimers to calculate Cα RMSDs of protB and assess its structural deviation relative to protA.

**c-e,** Structural comparison among the dimeric APLNR–$G_i$ complexes aligned on protA (**c**). Comparison of protB (**d**) reveals differences in the extent of its pre-dissociation from protA. The $G_i$ heterotrimer (**e**) undergoes a maximal rotation of 27°, as measured by αN helix.*The reported structures, including ELA–$G_i$ (PDB: 7W0N), cmpd644–$G_i$ (PDB: 7W0L), apelin–$G_i$ (PDB: 8XQE), JN241 (PDB: 8XQJ). All other structures were resolved in this study.

**a**

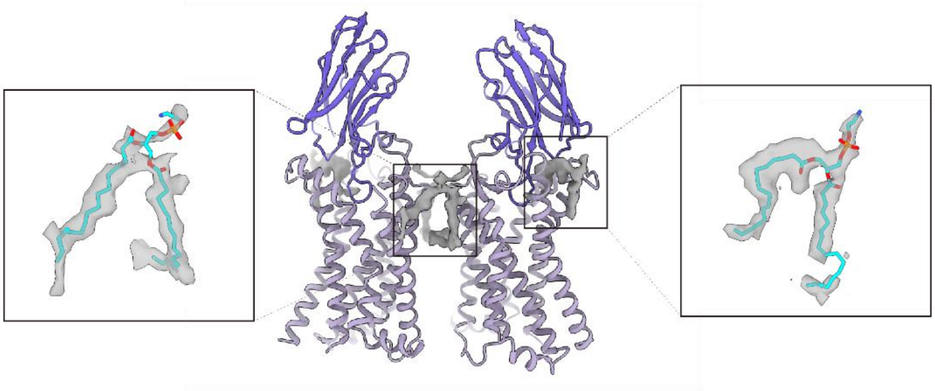

**b**

| | MβCD (mM) | 0 | 0.5 | 1 | 1.5 | 2 | 2.5 | 3 | 3.5 | 4 |
|---|---|---|---|---|---|---|---|---|---|---|
| Expression (%) | WT | 100.0±1.0 | 100.2±0.4 | 100.4±0.7 | 94.2±0.9 | 94.4±1.3 | 97.8±0.8 | 94.4±1.0 | 95.3±1.3 | 99.2±1.5 |
| | F101$^{3.24}$A | 87.0±1.4 | 88.1±0.9 | 88.3±1.3 | 80.9±1.5 | 79.8±1.5 | 82.8±1.3 | 82.1±1.6 | 81.5±1.7 | 86.4±1.7 |

**C**

| | WT | F101$^{3.24}$A | F39$^{1.43}$A | W51$^{1.55}$A | F54$^{1.58}$A | I66$^{2.41}$A | V73$^{2.48}$G | T87$^{2.62}$A | M162$^{4.58}$A |
|---|---|---|---|---|---|---|---|---|---|
| Expression (%) | 100.0±1.1 | 96.0±1.4 | 89.6±3.2 | 95.5±1.2 | 97.2±1.2 | 93.7±1.4 | 84.0±1.9 | 90.3±1.9 | 86.9±1.0 |

**Supplementary Figure 8. Characterization of phospholipids and analysis of cell surface expression in dimer interface mutations.**

**a,** Density map corresponding to the putative phospholipids.

**b, c,** Relative surface expression levels of mutants with (**b**) and without (**c**) MβCD treatment were measured by ELISA assay and normalized to the expression levels of WT.

**Supplementary Table 1. Cryo-EM data collection, model refinement and validation statistics of APLNR dimeric structures without G-protein.**

| | Apo (PDB: 9LQT) | JN241-9 LBS1a (PDB: 9LQU) | JN241-9 LBS1b (PDB: 9LQW) | JN241-9 LBS1c (PDB: 9LQX) | AMG986 LBS2a (PDB: 9LQY) | AMG986 LBS2b (PDB: 9LQZ) |
|---|---|---|---|---|---|---|
| **Data collection and processing** | | | | | | |
| Magnification | 150,540 | 150,540 | 150,540 | 150,540 | 150,540 | 150,540 |
| Voltage (kV) | 300 | 300 | 300 | 300 | 300 | 300 |
| Electron exposure ($e^{–}/Å^{2}$) | 52 | 52 | 52 | 52 | 52 | 52 |
| Defocus range (μm) | -1.0 ~ -2.5 | -1.0 ~ -2.5 | -1.0 ~ -2.5 | -1.0 ~ -2.5 | -1.0 ~ -2.5 | -1.0 ~ -2.5 |
| Pixel size (Å) | 0.93 | 0.93 | 0.93 | 0.93 | 0.93 | 0.93 |
| Symmetry imposed | C2 | C2 | C2 | C1 | C2 | C1 |
| Initial particle projections (no.) | 13,611,739 | 13,193,854 | 13,193,854 | 13,193,854 | 14,104,471 | 14,104,471 |
| Final particle projections (no.) | 265,860 | 97,020 | 101,362 | 138,605 | 306,883 | 189,970 |
| Map resolution (Å) | 2.7 | 2.7 | 2.7 | 2.7 | 2.4 | 2.5 |
| FSC threshold | 0.143 | 0.143 | 0.143 | 0.143 | 0.143 | 0.143 |
| Map resolution range (Å) | 2.0~3.0 | 2.0~3.0 | 2.0~3.2 | 2.2~3.0 | 2.0~3.0 | 2.0~3.0 |
| **Refinement** | | | | | | |
| Initial model used | 6KNM | 6KNM | 6KNM | 6KNM | 6KNM | 6KNM |
| Model resolution (Å) | 2.9 | 2.8 | 2.8 | 3.0 | 2.5 | 2.7 |
| FSC threshold | 0.5 | 0.5 | 0.5 | 0.5 | 0.5 | 0.5 |
| Model resolution range (Å) | 2.0~3.0 | 2.0~3.0 | 2.0~3.2 | 2.2~3.0 | 2.0~3.0 | 2.0~3.0 |
| Map sharpening *B* factor ($Å^{2}$) | -82.8902 | -77.9557 | -74.4693 | -84.909 | -66.4432 | -51.5988 |
| Model composition | | | | | | |
| Non-hydrogen atoms | 5,052 | 7,076 | 6,872 | 6,920 | 5,274 | 5,309 |
| Protein residues | 576 | 828 | 818 | 823 | 584 | 589 |
| *B* factors ($Å^{2}$) | | | | | | |
| Protein | 53.48 | 36.13 | 70.27 | 56.11 | 30.20 | 57.16 |
| Ligand | 70.27 | 36.50 | 69.45 | 41.83 | 39.68 | 66.75 |
| R.m.s. deviations | | | | | | |
| Bond lengths (Å) | 0.004 | 0.004 | 0.004 | 0.004 | 0.006 | 0.004 |
| Bond angles (°) | 0.815 | 1.004 | 1.037 | 0.983 | 1.065 | 1.009 |
| Validation | | | | | | |
| MolProbity score | 1.85 | 1.91 | 1.88 | 1.89 | 1.96 | 1.91 |
| Clashscore | 14.73 | 15.41 | 14.81 | 15.00 | 14.81 | 15.88 |
| Rotamer outliers (%) | 0.00 | 0.49 | 0.50 | 0.60 | 0.00 | 0.00 |
| Ramachandran plot | | | | | | |
| Favored (%) | 97.01 | 96.57 | 96.76 | 96.66 | 95.83 | 96.73 |
| Allowed (%) | 2.99 | 2.94 | 2.74 | 2.72 | 4.17 | 3.27 |
| Disallowed (%) | 0.00 | 0.49 | 0.50 | 0.62 | 0.00 | 0.00 |

### Supplementary Table 2. Cryo-EM data collection, model refinement and validation statistics of APLNR–G-protein complexes.

| | JN241-9 monomer (PDB: 9LR1) | JN241-9 dimer (PDB: 9LR2) | AMG986 monomer (PDB: 9LR3) | AMG986 dimer ACT2a (PDB: 9VPM) | AMG986 dimer ACT2b (PDB: 9VPN) |
|---|---|---|---|---|---|
| **Data collection and processing** | | | | | |
| Magnification | 150,540 | 150,540 | 150,540 | 150,540 | 150,540 |
| Voltage (kV) | 300 | 300 | 300 | 300 | 300 |
| Electron exposure ($e^{-}/Å^2$) | 52 | 52 | 52 | 52 | 52 |
| Defocus range (μm) | -1.0 ~ -2.5 | -1.0 ~ -2.5 | -1.0 ~ -2.5 | -1.0 ~ -2.5 | -1.0 ~ -2.5 |
| Pixel size (Å) | 0.93 | 0.93 | 0.93 | 0.93 | 0.93 |
| Symmetry imposed | C1 | C1 | C1 | C1 | C1 |
| Initial particle projections (no.) | 16,153,222 | 16,153,222 | 6,477,441 | 6,477,441 | 6,477,441 |
| Final particle projections (no.) | 187,827 | 198,168 | 63,229 | 64,609 | 95,469 |
| Map resolution (Å) | 2.5 | 3.0 | 2.8 | 4.0 | 4.0 |
| FSC threshold | 0.143 | 0.143 | 0.143 | 0.143 | 0.143 |
| Map resolution range (Å) | 2.0~3.5 | 2.0~4.0 | 2.0~3.2 | 3.0~5.0 | 3.0~5.0 |
| **Refinement** | | | | | |
| Initial model used | 8XZH | 8XZH | 8XZH | 8XZH | 8XZH |
| Model resolution (Å) | 2.5 | 3.5 | 2.8 | 6.5 | 6.3 |
| FSC threshold | 0.5 | 0.5 | 0.5 | 0.5 | 0.5 |
| Model resolution range (Å) | 2.0~3.5 | 2.0~4.0 | 2.0~3.2 | 5.0~7.0 | 5.0~7.0 |
| Map sharpening *B* factor ($Å^2$) | -51.5857 | -41.0765 | -46.165 | | |
| Model composition | | | | | |
| Non-hydrogen atoms | 9,913 | 13,115 | 9,046 | 8,586 | 5,769 |
| Protein residues | 1,271 | 1,671 | 1,149 | 1,423 | 1,442 |
| *B* factors ($Å^2$) | | | | | |
| Protein | 26.10 | 142.69 | 61.64 | 58.98 | 54.43 |
| Ligand | | 123.56 | 81.81 | | |
| R.m.s. deviations | | | | | |
| Bond lengths (Å) | 0.004 | 0.008 | 0.004 | 0.010 | 0.010 |
| Bond angles (°) | 0.947 | 1.170 | 0.987 | 1.773 | 1.748 |
| Validation | | | | | |
| MolProbity score | 1.51 | 1.83 | 1.53 | 0.71 | 0.71 |
| Clashscore | 6.21 | 11.41 | 8.98 | 0.28 | 0.27 |
| Rotamer outliers (%) | 0.37 | 0.54 | 0.10 | 0.00 | 0.00 |
| Ramachandran plot | | | | | |
| Favored (%) | 97.04 | 96.22 | 97.79 | 97.50 | 97.46 |
| Allowed (%) | 2.80 | 3.41 | 2.21 | 2.36 | 2.39 |
| Disallowed (%) | 0.16 | 0.37 | 0.00 | 0.14 | 0.14 |
